# Afrothismiaceae (Dioscoreales), a new fully mycoheterotrophic family endemic to tropical Africa

**DOI:** 10.1101/2023.01.10.523343

**Authors:** Martin Cheek, Marybel Soto Gomez, Sean W. Graham, Paula J. Rudall

## Abstract

*Afrothismia* is a genus of non-photosynthetic mycoheterotrophs from the forests of continental tropical Africa. Multiple phylogenetic inferences using molecular data recover the genus as sister to a clade comprising mycoheterotrophic Thismiaceae and the photosynthetic family Taccaceae, contrary to earlier placements of *Afrothismia* and Thismiaceae within Burmanniaceae. Morphological support for separating *Afrothismia* from the rest of Thismiaceae has depended on the zygomorphic flowers of *Afrothismia* (although some South American species of *Thismia* are also zygomorphic) and their clusters of root tubers, each with a terminal rootlet. The number of described species of *Afrothismia* has recently increased substantially, from four to 16, which has provided additional morphological characters that support its distinction from Thismiaceae. Most notably, the ovary in *Afrothismia* has a single stalked placenta, and circumscissile fruits from which seeds are exserted by placental elevation (in Thismiaceae, in contrast, there are three placentas, a deliquescing fruit lid, and the seeds are not exserted). *Afrothismia* stamens are inserted in the lower perianth tube where they are attached to the stigma, and individual flowers are subtended by a single large dorsal bract (in Thismiaceae, stamens are inserted at the mouth of the tube, free of and distant from the stigma, and each flower is subtended by a loose whorl of (2-)3(−4) bracts). Here we formally characterise Afrothismiaceae and review what is known of its development, seed germination, interactions with mycorrhizal Glomeromycota, biogeography, phylogeny and pollination biology. All but one (*Afrothismia insignis*; Vulnerable) of the 13 species assessed on the IUCN Red List of Threatened Species are either Endangered or Critically Endangered; one species (*A. pachyantha* Schltr.) is considered to be extinct.

## Introduction

The non-photosynthetic genus *Afrothismia* Schltr. (Schlechter 1906) is currently placed in the achlorophyllous tribe Thismieae (Burmanniaceae), together with *Thismia* Griff. (Griffith 1845), *Haplothismia* Airy Shaw (Airy Shaw 1952), *Oxygyne* Schltr. (Schlechter 1906), and the more recently described *Tiputinia* P.E. Berry & C.L. Woodw. (Woodward *et al*. 2007). Based on early molecular phylogenetic analyses that included plastid *rbcL, atpB* and nuclear 18S rDNA genes (Caddick *et al*., 2000a; 2000b), APGII (2003) and subsequent classifications combined Burmanniaceae *sensu stricto* with former Thismiaceae into a single family, Burmanniaceae *sensu lato*. However, the molecular-based analyses in these papers included contaminant plastid sequences (Lam *et al*., 2016), a common issue when using polymerase chain reaction (PCR) amplification to recover plastid DNA sequence data from mycoheterotrophic taxa. By contrast, morphological analysis (Caddick *et al*. 2002a) indicated the paraphyly of this group with respect to other Dioscoreales, based partly on the absence of septal nectaries in Thismiaceae (Caddick *et al*. 2000b; 2002a). In addition, more recent molecular phylogenetic data based in nuclear and mitochondrial sequences and whole plastid genomes (Merckx *et al*. 2009; 2010; Merckx & Smets 2014; Lam *et al*. 2016; 2018; Shepeleva *et al*. 2020; Lin *et al*. 2022) endorse earlier treatments (e.g., Jonker 1938; Maas van der Kamer 1998; APG I 1998) that support the separation of Thismiaceae from Burmanniaceae in different subclades of Dioscoreales, with Thismiaceae instead associated with Taccacaeae (e.g., Merckx *et al*. 2009; Merckx & Smets 2014). Moreover, studies based on mitochondrial and nuclear gene sets (Merckx *et al*. 2009; Merckx & Smets 2014; Merckx *et al*. 2017) have shown that Thismiaceae is itself paraphyletic, with *Afrothismia* recovered as the group of a clade comprising photosynthetic Taccaceae plus the rest of Thismiaceae. Strong support for this arrangement was recently recovered in phylogenomic analyses based on mitochondrial genomes (Lin *et al*. 2022). A general consensus has therefore emerged that *Afrothismia* is a phylogenetic lineage that is distinct from Burmanniaceae, Taccaceae and Thismiaceae (Fig. 1).

**Fig.1.**
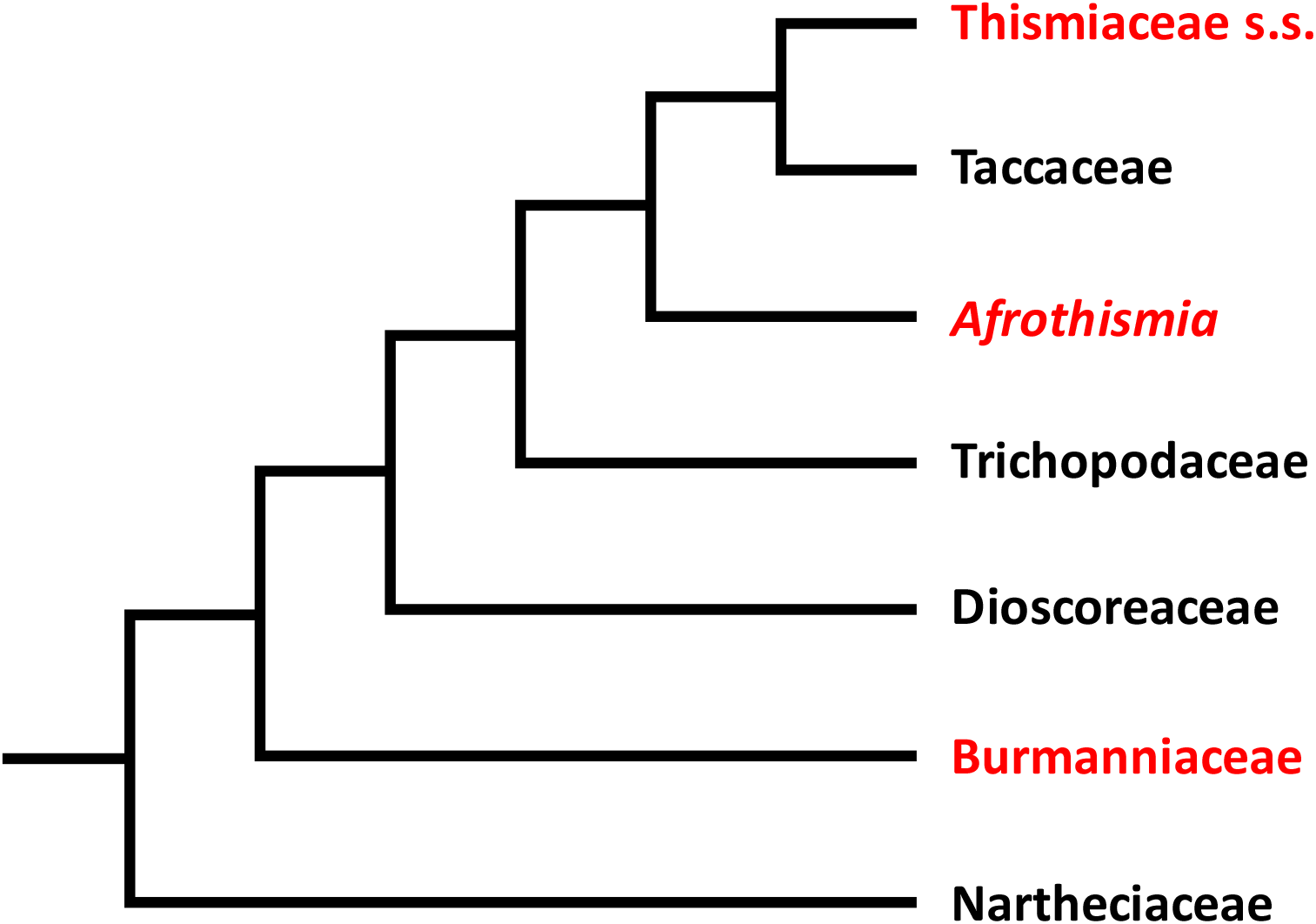
Higher-order relationships within Dioscoreales; the relationships follow Lin *et al*. (2022). Taxa in red are fully mycoheterotrophic (*Afrothismia*, Thismiaceae s.s.), or include both mycoheterotophic and photosynthetic taxa (Burmanniaceae).

To date, *Afrothismia* has been distinguished from *Thismia* by two morphological characters: zygomorphic flowers (but some South American species of *Thismia* are zygomorphic), and clusters of root tubers, each with a terminal rootlet (Maas van der Kamer 1998). However, in the last two decades, additional morphological data potentially support separate family status for *Afrothismia*, including new data associated with the description of numerous new species of *Afrothismia* (Cheek 2004a; 2007; 2009; Cheek *et al*. 2019; Cheek & Jannerup 2006; Dauby *et al*. 2008; Franke 2004; Franke *et al*. 2004; Maas-van de Kamer 2003; Sainge & Franke 2005; Sainge *et al*. 2005; 2013) and studies on ontogeny, floral morphology and mycorrhizal organization (Imhof & Sainge 2008; Imhof *et al*. 2020; Shepeleva *et al*. 2020). Here we review these recent studies, present evidence in support of family status for the genus, and formally describe and diagnose Afrothismiaceae as a new, fully mycoheterotrophic and monogeneric family in Dioscoreales.

Taccaceae Dumort., which is sometimes placed in or near Dioscoreaceae R.Br., is a pantropical family of 10–20 species, all placed in *Tacca* J.R. Forst. & G. Forst. Superficially these large, pantropical photosynthetic, terrestrial herbs seem unconnected with the non-photosynthetic Thismiaceae and Afrothismiaceae. However, as Rübsamen (1986) stated (translated from the German): “…Remarkable parallels exist in the structure of the anthers between Taccaceae and Burmanniaceae: the six stamens (of *Tacca*) look confusingly similar to those of *Afrothismia winkleri* or *Haplothismia*; the plate- or umbrella-shaped stigma (or the cap-like appearance of the style) is also reminiscent of the more bowl-shaped stigma of *Afrothismia*.”

## Methods

Nomenclature follows the Code (Turland *et al*. 2018). Authorship of names follows IPNI (continuously updated). The format of the description follows e.g. Cheek *et al*. (2019). Morphological terms follow Beentje & Cheek (2003). Herbarium codes follow Index Herbariorum (Thiers, continuously updated). All specimens cited have been seen. The conservation assessments cited follow the IUCN (2012) categories and criteria.

### Taxonomic Results

Diagnostic characters for *Afrothismia* are presented in Table 1 and Figs 1–4, including a comparison with Thismiaceae (*Haplothismia, Oxygyne, Thismia, Tiputinia*) and Taccaceae (*Tacca*).

**Table 1.** Diagnostic characters separating Thismiaceae and Taccaceae from Afrothismiaceae. Characters for Thismiaceae taken from Airy Shaw (1952), Caddick *et al*. (1998), Caddick *et al*. (2000a), Cheek *et al*. (2018a), Maas *et al*. (1998), Nuraliev *et al*. (2021), Woodward *et al*. (2007) and this paper (Fig. 3 D–I, J–L); those for Afrothismiaceae from Cheek (2004a; 2007; 2009), Cheek *et al*. (2019), Cheek & Jannerup (2006); Dauby *et al*. (2008), Franke (2004), Franke *et al*. (2004), Maas-van de Kamer (2003), Rübsamen (1986), Sainge & Franke (2005), Sainge *et al*. (2005; 2013), Imhof & Sainge (2008), Imhof *et al*. (2020) and this paper (Fig. 3 A–C); those for Taccaceae from Caddick *et al*. (1998), Caddick *et al*. (2000a), Drenth (1972), Kubitzki (1998), Watson & Dallwitz (1992 onwards).

|  | **Thismiaceae** (*Thismia*, Fig. 3D–I), *Haplothismia*, Fig. 3 J–L, *Oxygyne*, *Tiputinia*) | **Afrothismiaceae** (*Afrothismia*) (Fig. 2, 3A–C, 4) | **Taccaceae** (Fig. 5) |
| --- | --- | --- | --- |
| Rootstock | Rhizome or subglobose tuber | Clusters of globose bulbils, each with a terminal rootlet | Tuberous rhizome |
| Bracts | Loose whorl of (2–)3(–4) subequal bracts | A single subtending dorsal bract which covers the lower perianth tube (rarely bracts 2-3) | A single filiform bract subtending each flower |
| Inflorescence | Monochasial inflorescence of several flowers (thyrsoid with a single cyme). | Single or few-flowered, terminal or sympodial where more than one flower | Sympodial (cincinnus), contracted to an umbel |
| Floral symmetry | Actinomorphic (zygomorphic in *Thismia* species of former *Ophiomeris* Miers) | Zygomorphic, strongly (rarely only weakly e.g., *Afrothismia fungiformis*) | Actinomorphic (zygomorphic in aestivation) |
| Staminal insertion | At mouth of perianth tube | In lower perianth tube | In mouth or inside perianth tube |
| Stamen: stigma contiguity | Stamens sometimes basally fused into a ring around style; filamentous anther appendages not connected with stigma | Anthers becoming connected to the stigma surface at later stages via the apical connective appendage | Anthers curved beneath the style, not contiguous with the stigma; filaments short |
| Pollen apertures and surface sculpture | Mainly monoporate (monosulcate in *Haplothismia*, polyporate in some *Thismia*); sculpturing psilate or perforate | Monoporate; sculpturing coarsely reticulate | Monosulcate; finely reticulate, verrucate to slightly striate |
| Ovary | Inferior, unilocular with three parietal (often stipitate) placentas; septal nectaries absent | Inferior, unilocular with a single stalked, globose placenta; septal nectaries absent (Fig. 3A) | Inferior, unilocular with three parietal placentas; septal nectaries absent or rudimentary |
| Fruit | Opening by deliquescence (“withering”) of lid; seeds held inside the cup-like fruit | Circumscissile dehiscence of apex of fruit (pyxidium); seeds more or less exerted from fruit by placental elongation | A berry, rarely a loculicidal capsule |

### Taxonomic Treatment

**Afrothismiaceae** Cheek & Soto Gomez **fam. nov.**

Type genus: *Afrothismia* Schltr. (1906)

A single genus, *Afrothismia*, restricted to continental tropical Africa.

Description as for the genus (see below).

***Afrothismia*** Schltr. (Schlechter 1906; Jonker 1938; Cowley 1988; Maas-van de Kamer 1998; Cheek 2009)

Type of genus: *Afrothismia winkleri* (Engl.) Schltr. Jonker (1938: 223)

*Perennial non-photosynthetic mycoheterotrophic herbs* entirely lacking green tissue, with only the flower or fruit emerging above the leaf-litter. *Stem* (rhizome), colourless, opaque, succulent, concealed in substrate, spreading more or less horizontally, becoming vertical when flowering, sparsely branched or unbranched. *Scale-leaves* sparse, alternate, ovate-triangular, minute, axillary buds globose, minute. *Bulbil clusters* subglobose in outline, bulbils 15–40, each globose, with an apical rootlet (Fig. 4B).

*Inflorescence* 1–few-flowered, terminal or sympodial where more than 1 flower (Fig. 2B). *Flowers* strongly or weakly zygomorphic, subtended by a large dorsal colourless ovate bract (Fig. 2A, 3A, 3B, 4D). *Perianth tube* usually colour-patterned, translucent, white, red and/or purple, erect to horizontal, straight, S-shaped or angled, globose or subcylindrical, constricted or not, separated internally usually by an annulus, into two parts, upper and lower, outer surface smooth, ribbed or papillate; mouth of tube projecting beyond insertion of the lobes, ribbed or not, partly covered with an operculum (corona) or not, aperture orbicular, hemi-orbicular or elliptic; post-anthesis deliquescing. *Perianth lobes* six, yellow, white, red or purple, patent, forward-directed or curved, equal or unequal, triangular or filiform, entire or lacerate.

**Fig. 2.**
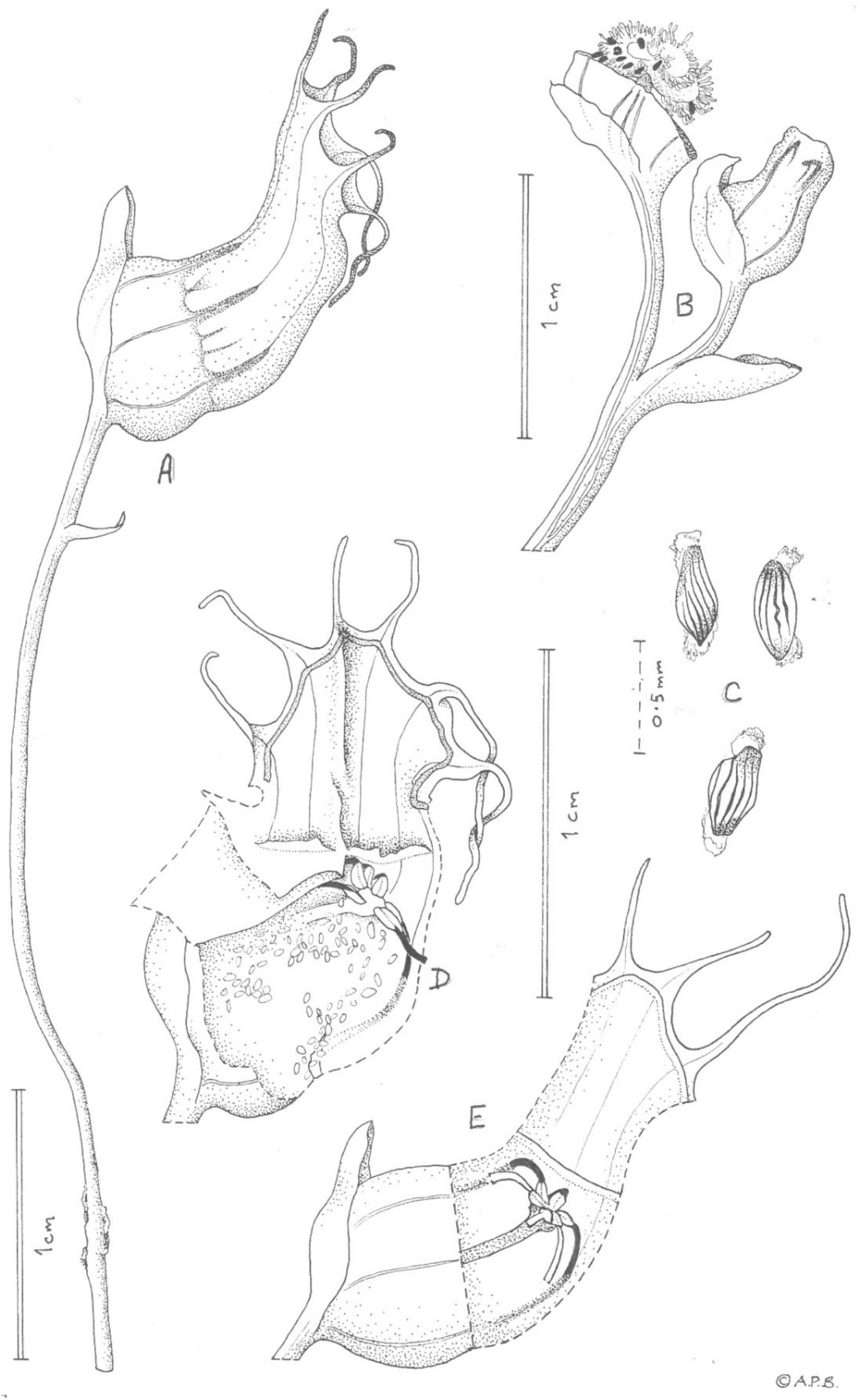
Afrothismia zambesiaca. **A** habit, flowering stem; **B** dehisced fruit (showing the placenta with its seeds, exserted by the ‘placentophore’) on sympodial inflorescence; **C** seeds, showing an appendage, a possible elaiosome at each end; **D** dissected flower; **E** flower in section, reconstructed from D. All drawn from *Exell, Mendonça & Wild* 1066 (holotype, K) by ANDREW BROWN.

**Fig. 3.**
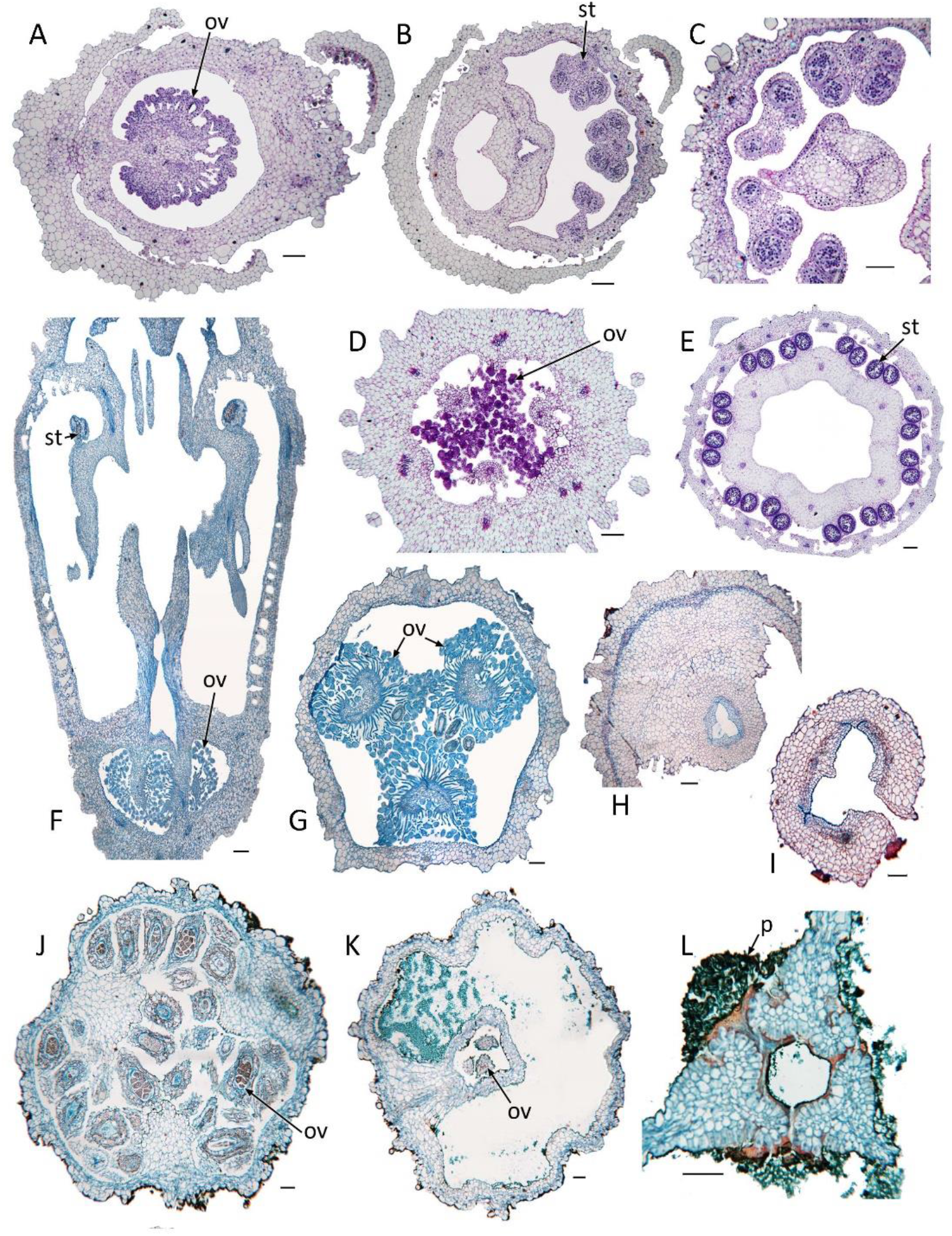
Photomicrographs of sections of flowers. **A–C** *Afrothismia hydra* Sainge & T.Franke, serial transverse sections of zygomorphic flower bud; **A** ovary with a single placenta on a long stalk (‘placentophore’); **B** top of ovary showing base of hollow style and stamens on one side; **C** hollow style. **D, E** *Thismia aseroe* Becc., serial transverse sections of flower bud; **D** ovary with three equal placentas and numerous ovules (ov) at early developmental stages; **E** stamen tube with external developing anthers. **F–I** *Thismia episcopalis* (Becc.) F.Muell.; **F** longitudinal section of young flower bud showing apical placentas in ovary and stamen connectives extending downwards towards stigma (not connected with stigma at this stage); **G–I serial** transverse sections of young flower bud; **G** three equal placentas in ovary; **H** top of ovary and style base; **I** hollow style. **J–L** *Haplothismia exannulata* Airy Shaw, serial transverse sections of flower; **J** three equal placentas in ovary; **K** top of ovary with mucilage accumulation around style base; **L** style with mass of germinating pollen tubes (p) inserted between carpel margins. Labels: b = bract, ov = ovule, p = pollen-tube mass, st = stamen. Scales = 100 μm.

**Fig. 4.**
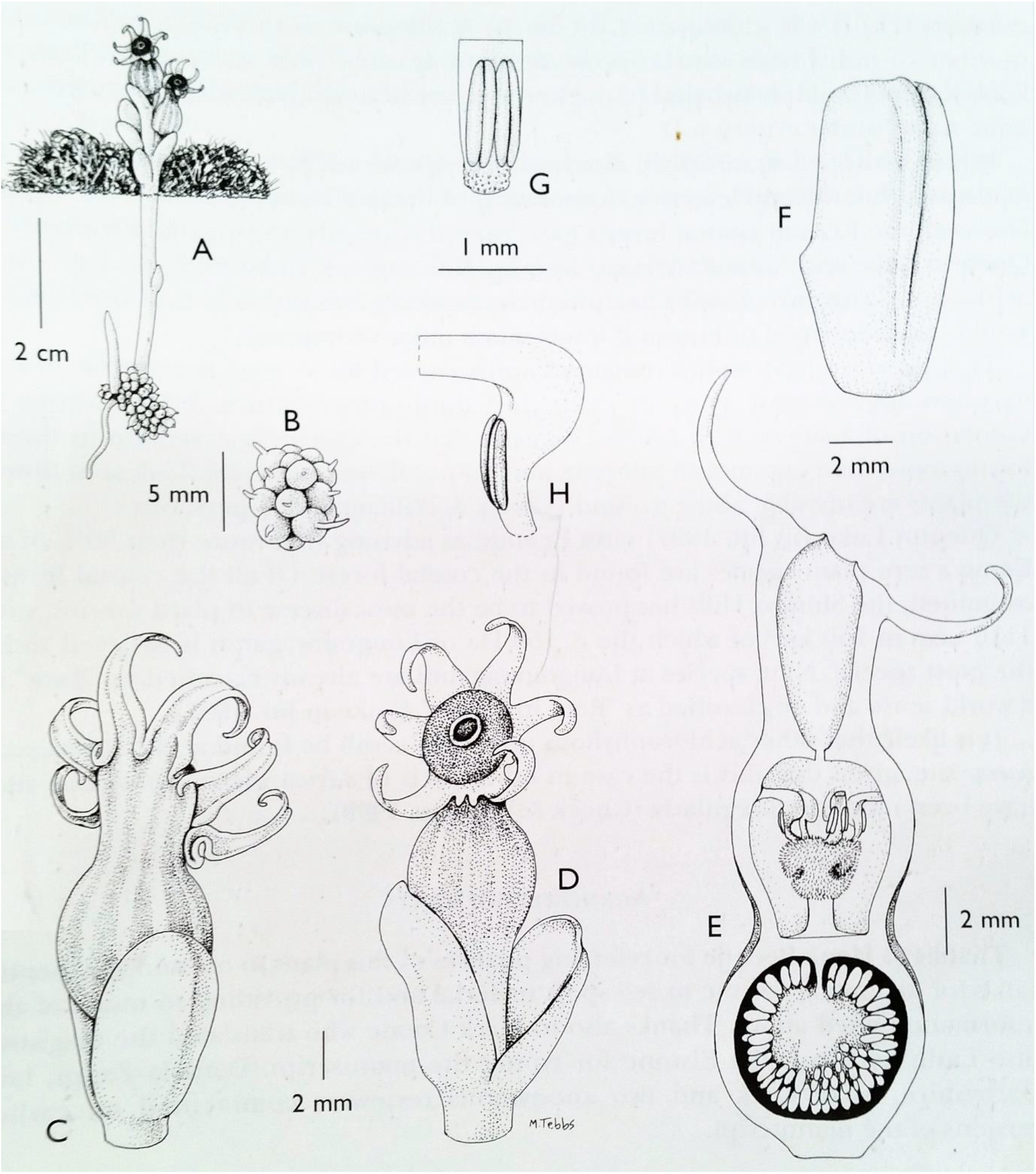
Afrothismia baerae. **A** habit, flowering stem; **B** cluster of root tubers, each with terminal root; **C** flower with bract (rear view); **D** flower, frontal view; **E** longitudinal section of flower (non-median); **F** flower-subtending bract; **G** detail of stamen and connective; **H** side view of the free part of the stamen showing the expanded filament-connective structure above the anther cells. All drawn from *Baer* s.n. (K) by MARGARET TEBBS.

**Fig. 5.**
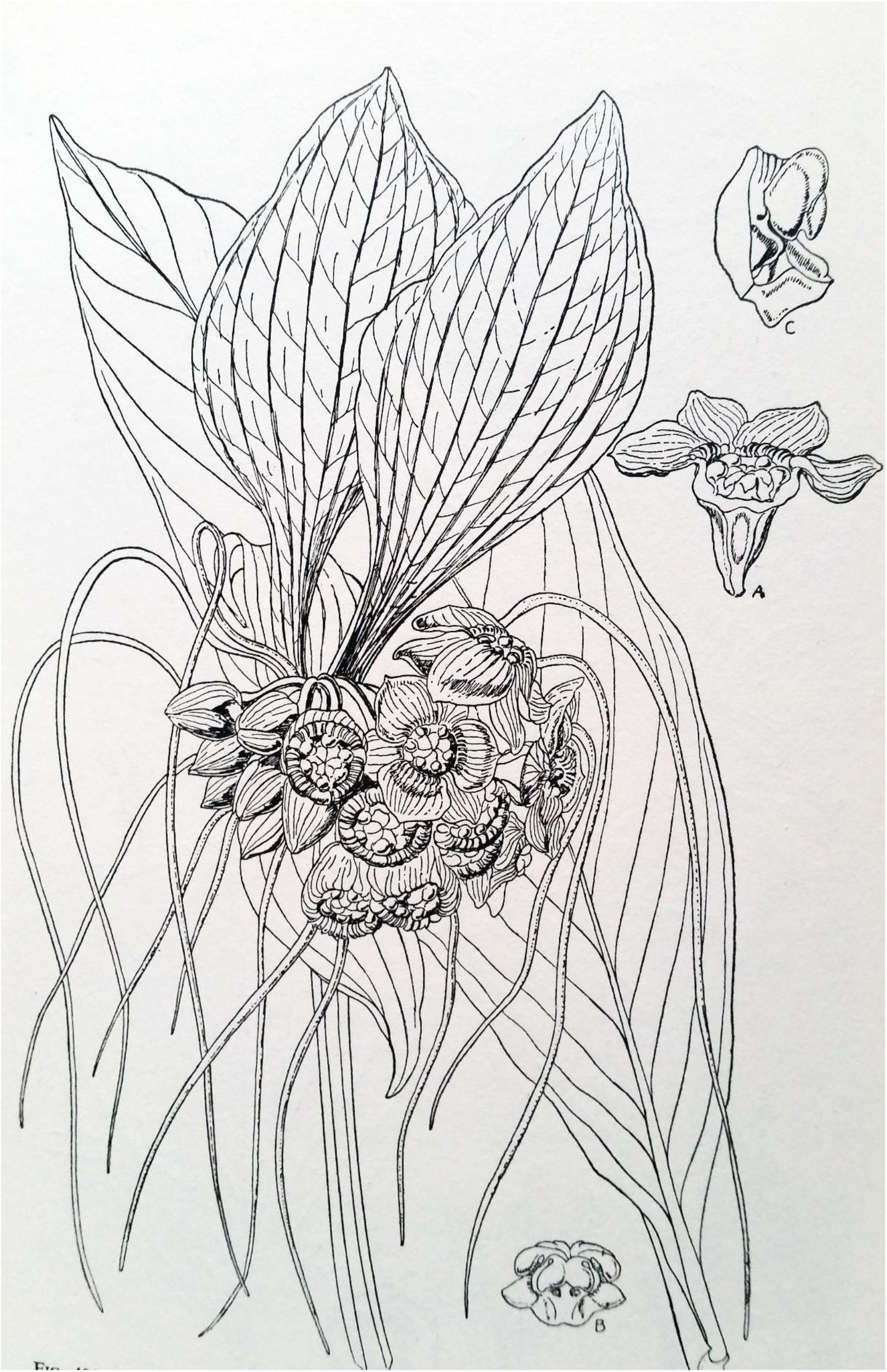
Tacca cristata. **A** flower; **B** stamens, showing contiguity with stigma; **C** one stamen from the side showing the filament-connective hood; **D** inflorescence and leaf-blade. From Carter (1962).

*Stamens* 6, inserted on distal part of lower perianth tube (epipetalous) below the annulus, staminal filaments dorsiventrally slightly flattened or clavate, the apex and basal part of the connective swollen and sometimes geniculate (Fig. 4H), arching inward and downward to stigma, glabrous, papillate or hairy; anther thecae two, elliptic, introrse, separated by and often embedded in the connective, distal connective appendage papillate (Fig. 4G), firmly adnate to stigmatic surface (Fig. 4E), pollen monoporate, surface reticulate.

*Ovary* inferior, campanulate, unilocular, placentation axile, placenta globose sometimes 3-lobed, massive, attached at base and apex by cylindrical stalks; ovules numerous, anatropous often on long funicles (Fig. 3A). Style cylindrical, short, hollow (Fig. 3B, C); stigma obconical thick, usually slightly 6-lobed or not, surface densely papillate or hairy (Fig. 4E).

*Fruit* campanulate, dehiscence circumscissile, the lid of the fruit (perianth floor) detaching completely, exposing the seed-covered placenta which is projected more or less completely above or far above the fruit wall by the elongation of the placental base or placentophore. *Seeds* numerous, narrowly obovoid or ellipsoid, reticulate, lacking appendages or with a swollen structure (possible elaiosomes) at each end. (Fig. 2).

#### RECOGNITION (diagnosis)

Afrothismiaceae fam. nov. differing from Thismiaceae Aghardh in that the ovary has a single stalked, globose placenta, fruits circumscissile (pyxidium), seeds exserted from fruit by placental elevation (vs. parietal, fruit lid deliquescing, seeds included), rootstocks with clusters of globose root tubers, each with a terminal root (vs. a single tuber or roots fleshy), stamens inserted in mid or lower part of perianth tube, below the annulus; anther appendages connected to stigma at anthesis (vs. inserted near mouth, distant and free from stigma).

#### DISTRIBUTION & BIOGEOGRAPHY

Continental tropical Africa, from Nigeria east to Kenya, south to Malawi. Of the 16 described species, 12 occur in Cameroon, one of these extending west to Nigeria, another East to Gabon. Gabon has one described endemic species, Uganda another (currently treated at varietal rank), Kenya and Malawi both have one, and Tanzania two. However, several undescribed species are known to have been collected in Cameroon (Sainge *et al*. 2017), Gabon (MC, pers. obs) and one in Tanzania (*Afrothismia “arachnites”*, Rübsamen (1986)). In Cameroon, the species are concentrated in the Cross-Sanaga Interval (Cheek et al 2001), where nine of the 12 described Cameroon species occur, seven of which are endemic. This area contains the highest vascular plant species and generic diversity per degree square in tropical Africa, with endemic genera such as *Medusandra* Brenan (Peridiscaceae, Barthlott *et al*. 1996; Dagallier *et al*. 2020; Soltis *et al*. 2007; Breteler *et al*. 2015). The species of *Afrothismia* in this area show the full range of floral and root morphology documented in the genus. The genus is unrecorded from the Congo basin and there is a c. 2200 km disjunction between the species of Lower Guinea (West-Central Africa) and the westernmost East African record in Uganda.

The single most species-diverse area for *Afrothismia* is Mt Kupe in S.W. Region, Cameroon, within the Cross-Sanaga Interval. Here five species have been recorded: *A. saingei* (Franke 2004), *A. fungiformis* (Sainge *et al*. 2013), *A. winkleri* (Cheek *et al*. 2004), *A. hydra* (Onana & Cheek 2011) and *A. kupensis* (Cheek *et al*. 2019). The first and last species are thought to be endemic to Mt Kupe. The Mt Kupe area has been the source of numerous other new species to science e.g. (Stoffelen *et al*. 1997; Cheek & Csiba 2002; Cheek 2003) and even new genera, including *Kupeantha* Cheek (Rubiaceae, Cheek *et al*. 2018b) and (another non-photosynthetic mycoheterotroph) *Kupea* Cheek & S.A. Williams (Triuridaceae, Cheek *et al*. 2003) subsequently found to extend to East Africa (Cheek 2004b).

At Mt Kupe, *Afrothismia* species occur with other achlorophyllous mycoheterophic plant species in the families Gentianaceae, Burmanniaceae and Triuridaceae, including one observed site of c. 20 m x 20 m with six mycoheterotrophic species, including *Afrothismia kupensis* (Cheek & Williams 1999; Cheek 2006), equalling the record in the other most species-diverse site ever recorded in Africa, at Moliwe, Mt Cameroon (Cheek & Ndam 1996, analysing collection records from Schlechter), which is now converted into agricultural plantations.

The first records of several of the mycoheterophs at Mt Kupe were made in the course of intensive botanical surveys conducted over several seasons, including non-photosynthetic mycoheterophic plant specialists, to support conservation management. The surveys usually resulted in a botanical conservation checklist (in the case of Mt Kupe, Cheek *et al*. 2004). However similar surveys at several other locations in Cameroon with lowland or submontane forest that might be expected to host *Afrothismia* failed to uncover any plants, even in the appropriate late wet season (Cheek *et al*. 2000; 2010; 2011; Harvey *et al*. 2004; 2010). These records suggest that *Afrothismia* species are not ubiquitous in any apparently suitable habitat, but genuinely localised and rare, even in Cameroon, where the majority of the known species are recorded.

#### HABITAT

Lowland and submontane evergreen forest; c. 200-1150 m altitude.

#### ETYMOLOGY

Taken to signify “African *Thismia*”.

#### CONSERVATION STATUS

Of the 17 published taxa (16 species) of *Afrothismia*, 14 have been assessed for their IUCN extinction risk status (Table 2). One has been assessed as Vulnerable, two Endangered and 11 of the 14 are assessed as Critically Endangered (CR), the highest category of threatened status before extinction. The level of CR species in the *Afrothismia* (c.78%) is exceptionally high and exceeds that of extremely threatened genera of comparable size e.g. *Inversodicraea* (Podostemaceae) with 14/30 (c.46%) CR species (Cheek *et al*. 2017). *Afrothismia pachyantha* has subsequently been declared extinct (Cheek *et al*. 2019), based on multiple attempts to relocate non-photosynthetic mycoheterotrophic plants at its sole locality by teams of mycotroph specialists since 1991. Its habitat has been cleared for plantation and small-holder agriculture, including rubber (*Hevea brasiliensis* ([Willd](https://www.ipni.org/a/11692-1). ex [A.Juss](https://www.ipni.org/a/4593-1).) [Müll.Arg](https://www.ipni.org/a/23712-1).).

**Table 2.** Published IUCN species extinction risk assessments for species of *Afrothismia*. All but three taxa (two species) have been assessed and can be viewed on iucnredlist.org. IUCN (2012) codes: NE = Not Evaluated; LC = Least Concern; NT = Near Threatened; VU = Vulnerable; EN = Endangered; CR = Critically Endangered. *Afrothismia pachyantha* has subsequently been discovered to be extinct (Cheek *et al*. 2019) but the IUCN assessment remains to be updated.

| Species | Reference | NE | LC | NT | VU | EN | CR |
| --- | --- | --- | --- | --- | --- | --- | --- |
| *A.amietii* | Cheek (2018a) |  |  |  |  |  | X |
| *A.baerae* | Eastern Arc Mountains & Coastal Forests<br>CEPF Plant Assessment Project (2009a) |  |  |  |  |  | X |
| *A.foertheriana* | Cheek (2018b) |  |  |  |  |  | X |
| *A.fungiformis* | Whittaker & Cheek (2021) |  |  |  |  | X |  |
| *A.gabonensis* |  | X |  |  |  |  |  |
| *A.gesnerioides* | Cheek (2018c) |  |  |  |  |  | X |
| *A. hydra* | Cheek & Lovell (2020) |  |  |  |  | X |  |
| *A.insignis* | Eastern Arc Mountains & Coastal Forests<br>CEPF Plant Assessment Project (2009b) |  |  |  | X |  |  |
| *A.korupensis* | Cheek (2017) |  |  |  |  |  | X |
| *A.kupensis* | Cheek & Rotton (2022) |  |  |  |  |  | X |
| *A.mhoroana* | Mwangoka *et al.* (2021) |  |  |  |  |  | X |
| *A.pachyantha* | Cheek (2004c) |  |  |  |  |  | X |
| *A.pusilla* | Harvey & Cheek (2021) |  |  |  |  |  | X |
| *A.saingei* | Cheek & Rokni (2017) |  |  |  |  |  | X |
| *A.winkleri* | IUCN SSC East African Plants Red List<br>Authority (2013) |  |  |  |  |  | X |
| *A.winkleri* var.<br>*budongensis* |  | X |  |  |  |  |  |
| *A.zambesiaca* |  | X |  |  |  |  |  |
| Total |  | 3 | 0 | 0 | 1 | 2 | 11 |

Other locations lost due to agricultural clearance include the type location of *Afrothismia winkleri* at Muea (Onana & Cheek 2011). This represents the main threat faced by *Afrothismia* species; they are acutely susceptible because most occur in either a single or only two to three locations, and at each location, their population may occupy only 2–10 m^2^. It is expected that additional species will shortly become extinct, if they are not already so. The forest at Mt Kala, type and sole locality for *A. amietii* and *A. pusilla*, is being cleared for housing development. *Afrothismia baerae* has not been seen for 20 years since it was first collected in 2002, despite annual monitoring (Quentin Luke *pers. comm*. to MC 2022). However, other species not seen for decades, such as *A. zambesiaca*, unseen since the type gathering in 1955, cannot be assumed extinct because there have been no targeted efforts to refind them. In order to support their protection, in 2022, almost all Cameroon *Afrothismia* species were included within a network of Important Plant Areas (IPAs or TIPAs, Darbyshire *et al*. 2017; https://www.kew.org/science/our-science/projects/tropical-important-plant-areas-cameroon), however, these species lack legal protection. Eight *Afrothismia* species are included in the Cameroon Red Data Book (Onana & Cheek 2011: 353-356).

Non-photosynthetic mycoheterotrophic plants such as *Afrothismia* have never been recorded as being successfully cultivated. We do not yet know which autotrophic plant species their soil fungal symbionts depend upon. Presumably, to succeed in growing *Afrothismia* from seed, one would need to have both the required species of fungal symbiont and suitable autotrophic plant partner(s) already established. There is no record that this goal has either been achieved or attempted. It is likely that *Afrothismia* might have orthodox seeds since they are small and dry, but no species are known to be seed-banked, and seed banking is of little value if the seeds cannot be grown to produce viable plants.

The IUCN convention is that species are not considered for inclusion on their website (iucnredlist.org) until they are formally published. This makes it more urgent to publish species so that they can be formally Red Listed and afforded a higher level of protection than they would otherwise obtain (Cheek *et al*. 2020). Therefore, publication of the seven known but still undescribed species of *Afrothismia* remains a priority.

### Typification

In the generic protologue, two species were published (Schechter 1906). Jonker effectively chose one of these as the lectotype of the genus by giving *Afrothismia winkleri* as “Type species” (Jonker 1938: 223).

### Species discovery

*Afrothismia winkleri* was first published as *Thismia winkleri* (Engler 1905) before becoming the foundation of Schlechter’s genus *Afrothismia* Schltr., joined by *A. pachyantha* Schltr. (Schlechter 1906). Both species were first collected on the slopes of Mt Cameroon. The third species was collected in the Usambara Mts, now in Tanzania in East Africa, by Peter (s.n., B), who labelled it as *Afrothismia arachnites* n. sp., but never formally published it. Until today, this species seems to have been overlooked, except by Rübsamen (1986). Two new taxa were added to the genus more than 80 years later by Cowley (1988), *Afrothismia winkleri* var. *budongensis* Cowley (Uganda) and *A. insignis* Cowley (Tanzania). By the end of the 20^th^ Century, *Afrothismia* was known to have only three species, two of which occurred at Mt Cameroon, one endemic (Cheek & Ndam 1996). A large increase in species discovery and publication, arising mainly from Cameroon, began 15 years after Cowley (1988). *Afrothismia gesnerioides* H.Maas (Maas-van der Kamer 2003) of Cameroon, was soon followed by *Afrothismia baerae* Cheek (2004) of Kenya, and four species from S.W. Region, Cameroon: *A. saingei* T.Franke (2004), *A. foertheriana* T. Franke *et al*. (Franke *et al*. 2004), *A. hydra* Sainge & T.Franke and *A. korupensis* Sainge & T.Franke (Sainge *et al*. 2005; Sainge & Franke 2005). *Afrothismia mhoroana* Cheek of Tanzania (Cheek & Jannerup 2006), *A. amietii* Cheek (2007) of Cameroon, *A. gabonensis* Dauby & Stévart (Dauby *et al*. 2008) of Gabon, and *A. zambesiaca* Cheek (2009) of Malawi. The most recently published taxa are three species from Cameroon: *Afrothismia pusilla* Sainge & Kenfack, *A. fungiformis* Sainge & Kenfack (both Sainge *et al*. 2013), *A. kupensis* Cheek & S.A. Williams (Cheek *et al*. 2019).

Sainge *et al*. (2017) mentioned three further still undescribed species (referred to as sp. a, b, d, respectively) that he had collected in Cameroon and a fourth that he had seen from Gabon, which he termed sp. c (*Boupouya et al*. 674). The first author has seen photos of two additional undescribed species from the Cristal Mts of Gabon, *Bidault* 5044 and *Bidault* 5478. Additionally, there is *Peter s.n*. (*Afrothismia* “arachnites”) of Tanzania (Rübsamen 1986). Thus, seven undescribed species remain to be published, which would take the total number of species in the genus and family to 23, exceeding the 20 species currently accepted in Taccaceae (Plants of the World online, continuously updated).

Taking existing point data records for *Afrothismia* and using Maxent and an ecological niche modelling approach to map the potential range, Sainge *et al*. (2017) predicted highly suitable areas where additional species might be found in Sierra Leone, Liberia, Ivory Coast, Nigeria, Cameroon, Equatorial Guinea, Gabon, Republic of Congo, and Democratic Republic of Congo. However, regarding the first three countries, which equate to Upper Guinea, no *Afrothismia* species have been found to date, although other recent discoveries of achlorophyllous mycoheterophic plant species have been made there by specialists (e.g. Cheek & van der Burgt 2010).

### Mycorrhizal relationships in *Afrothismia*

Combined molecular results from 18S *r*DNA, ITS and *atp*A were used to study the phylogeny of several species of *Afrothisma* and their fungal partners from three locations in SW Region Cameroon (Merckx & Bidartondo 2008). The species of *Afrothismia* included were *A. winkleri, A. hydra, A. foertheriana, A. kupensis* (as *A. gesnerioides*) and *A. korupensis*. All the fungal symbionts were placed in the *Glomus* sp. A lineage of the Glomeromycota, and there was no fungal lineage overlap among the different species of *Afrothismia*. No other fungi or fungal-like organisms were identified, apart from a stramenopile (non-mycorrhizal, assumed pathogen) in *A. foetheriana*. Franke *et al*. (2006), investigating the symbionts of seven taxa of *Afrothismia* also from S W Region, Cameroon, had also found them all to be exclusively *Glomus* sp A. lineage. However, Imhof (2006) reported an unidentified second fungal species in material of *A. gesnerioides*. A delayed co-speciation pattern between the plant species and the *Glomus* lineages was revealed (Merckx 2008; Merckx & Bidartondo 2008): the divergence time estimates for the *Glomus* nodes were older than for their corresponding nodes in *Afrothismia*.

Glomeromycota form vascular arbuscular mycorrhiza with about 90% of all land plants (Cheek *et al*. 2020). *Glomus* group A are also symbionts with Taccacaeae (Merckx 2008). There is evidence for the origin of Glomeromycota before 400 Mya (Strullu-Derrien *et al*. 2018).

Many other non-photosynthetic mycoheterophic plants also depend on Glomeromycota as symbionts, including Thismiaceae s.s., Burmanniaceae s.s., and Triuridaceae s.s. However non-photosynthetic mycoheterophic Ericaceae and some Orchidaceae are mainly ectomycorrhizal and depend on Ascomycota and Basidiomycota as symbionts.

### Dates of origin and diversification of *Afrothismia*

Using two different relaxed molecular clock models on the same study set as used above, the origin of *Afrothismia* was estimated as 91±11 Mya and 120± 11 Mya, with diversification of the genus starting around 50± 13 Mya and 78± 9 Mya (Merckx 2008; Merckx & Bidartondo 2008; Merckx *et al*. 2010; Merckx & Smets 2014; Merckx *et al*. 2017). *Afrothismia kupensis* diverged 50 Mya (its fungal symbiont 219 Mya), *A. korupensis* 34 Mya (its symbiont and that of *A. foetheriana* 122 Mya), while *A. hydra* and *A. winkleri* diverged from each other 0.8 Mya (their symbionts 66 Mya).

### Morphological trends and evolution: flowers

The species of *Afrothismia* included in the molecular phylogenetic analyses of Merckx & Bidartondo (2008) and Merckx *et al*. (2009) are representative of the range of variation in floral morphology currently known in *Afrothismia*. Here, we briefly describe these patterns, referring to other species that share similar morphology, accepting that this similarity might result in part from convergence rather than recent common ancestry. It is likely that these apparent morphological trends are associated with attracting pollinators (see below; pollination biology).

To date, no molecular phylogenetic analyses have included all 16 described species of *Afrothismia*. Merckx & Bidartondo (2008) and Merckx *et al*. (2009) included five and six species respectively, and Shepelova *et al*. (2020) included six species. In these analyses, *A. kupensis* (as *gesnerioides*) was sister to the remaining *Afrothismia* species. Both *A. gesnerioides* and *A. kupensis* share similar morphology and are probably closely related, both species possessing tepals that are triangular and non-filamentous and a horizontal perianth tube that is only slightly sinusoidal in the upper part. Among other species, *A. korupensis* shows the more-or-less filamentous perianth lobes that are present in all other species of the genus except *A. gesnerioides* and *A. kupensis*. In *A. korupensis*, the perianth tube is vertical and the proximal and the majority of the distal part aligned on the same axis. Only the mouth and uppermost part of the distal tube is orientated horizontally, with a projecting corona partly occluding the mouth. This pattern, of a vertical perianth tube with horizontal mouth, is also seen in *A. pusilla, A. fungiformis* (both also in Cameroon), *A. baerae* (Kenya) and *A. “arachnites”* (Tanzania). Another species, *A. foertheriana*, has a highly reduced distal perianth tube, the campanulate proximal tube comprising the majority and being about as wide as long, and the lobes and corona bear numerous short projections that are also seen in *A. pusilla. Afrothismia amietii* of Cameroon also shares this pattern, differing in the distal tube being entirely absent. The remaining species examined, *A. hydra* and *A. winkleri*, are notable for the proximal perianth tube being ± horizontal, the distal tube being angled vertically, forming an L-shape. This pattern is seen, with variations, throughout the range of the genus, occurring also in *A. gabonensis* (Gabon), *A. insignis, A. mhoroana* (both Tanzania) and *A. zambeziaca* (Malawi). It is also among these species that yellow is included among the flower colours. All other species are coloured in a combination of purple or dark dull red, usually with white (though white is absent from the perianth of *A. foertheriana*). The flowers of *Afrothismia mhoroana* are yellow and white in colour, lacking purple or dark red colouring entirely.

### Mycorrhizal trends and evolution

A series of studies of the mycorrhizal structures of *Afrothismia* (Imhof 1999; 2006; Imhof *et al*. 2013; 2020) also found signs of substantial ongoing evolutionary diversification. The studies involved *Afrothismia winkleri* (identification to be confirmed, based on *Wilks* 1179 of Gabon, Imhof 1999), *Afrothismia gesnerioides* (based on *de Winter* 91(L), S Region Cameroon, Imhof 2006), *Afrothismia kupensis* (as *A. gesnerioides*) and likely *Afrothismia winkleri* (as *Afrothismia saingei*) (Imhof *et al*. 2013) and *Afrothismia hydra, A. korupensis, A. gesnerioides* and likely *Afrothismia winkleri* (as *Afrothismia saingei*) (Imhof *et al*. 2020).

According to Imhof *et al*. (2020), the root-shoot combination of *Afrothismia winkleri* exhibits one of the most complex mycorrhizal colonization patterns described to date. It shows four different hyphal shapes (straight, looped, inflated coils, degenerating coils) in six separate tissue compartments (filiform root, root epidermis, third root layer, root cortex parenchyma, shoot cortex at root clusters, shoot cortex apart from root clusters). Interconnections between all hyphal shapes demonstrated that they belong to the same fungus. In addition, the long filiform roots of this species were interpreted as being especially efficient in facilitating penetration by fungi. In contrast, the mycorrhizal pattern in *A. gesnerioides* is comparatively simple, with three hyphal forms in five tissue compartments, and the short blunt roots interpreted as being less efficient. *Afrothismia hydra* and *A. korupensis* were found to be intermediate in complexity between the foregoing species.

Imhof *et al*. (2020) suggested that the differences between four *Afrothismia* species reflected a transitional change towards increasing functional complexity and strict partitioning of conveyance (straight hyphae) and storage purposes (inflated hyphal coils). They concluded that since investigations on the mycorrhizal structures of *Thismia* spp. describe a disparate and much less sophisticated colonization pattern than in *Afrothismia*, this difference supports taxonomic separation from Thismiaceae.

### Ontogeny

Caddick *et al*. (2000) described comparative floral ontogeny in several Dioscoreales, including two species of *Thismia* and four species of *Tacca*. Nuraliev *et al*. (2021) provided a detailed description of both flower and inflorescence ontogeny in several species of *Thismia*, noting variation in placentation in different species, including some in which placentation is parietal at the base and columnar above. Flower buds of *Afrothismia hydra, Haplothismia exannulata* and two species of *Thismia* are illustrated here (Fig. 3). Imhof & Sainge (2008) documented development in *Afrothismia hydra* from seed to seed-dispersal.

In *A. hydra*, seeds germinate with root tissue only, disrupting the seed coat and developing a primary ovoid root tubercle. The hypocotyl, cotyledons and shoot are not visible during germination. A second tubercle arises at the proximal end of the first one and subsequent root tubercles with filiform extensions develop sequentially, resulting in a small root aggregate. The root aggregate enlarges, forming a central axis to which all roots are connected. This axis has a growth pole where new root tubercles arise; it later develops into a stem with scale leaves, finally terminating in a flower. After anthesis, the corolla tube disintegrates, leaving a pyxidium which opens by means of a peculiar elongating placenta, which Imhof & Sainge (2008) termed a ‘placentophore’, also shown in our material of *A. hydra* (Fig. 3A). The placentophore later elevates the placenta with attached seeds above the flowering level and is interpreted as an adaptation to ombrohydrochory (rain-operated seed dispersal). The placentophore can extend to 4–5 times the length of the fruits and seems to be the result of meristem growth rather than cell elongation (Imhof & Sainge 2008).

### Pollen structure

In her excellent systematic exploration of the embryology, seeds, and pollen of Burmanniaceae s.l. and Corsiaceae, Rübsamen (1986) made an SEM study of the pollen and seeds of two species of *Afrothismia, A.winkleri* (based on *Zenker* 3613, B) and *A. “arachnites”* (based on *Peter s.n*., B), using dried herbarium material. Caddick *et al*. (1998) described microsporogenesis and pollen morphology in most genera of Dioscoreales, including *Thismia* and *Tacca*. In *Afrothismia*, pollen grains are released as single units, as in most Dioscoreales (rarely as tetrads in some Burmanniaceae and *Thismia* species); they are plano-convex, monoporate (ulcerate), with pollen lengths of the two species 24 μm and 14–16 μm, respectively (Rübsamen 1986). The surface sculpture is coarsely reticulate, the muri “caterpillar like”, occasionally with tiny pores, compared with more finely perforate (rarely smooth) in *Thismia* (Rübsamen 1986). Rübsamen (1986) stated that “…Because of the exine sculpture, the genus *Afrothismia* (with a reticulate exine surface) is also clearly separated from the genus *Thismia*.“

### Pollination biology

Within *Afrothismia*, pollination has been recorded only in *A. kupensis* (Cheek *et al*. 2019). Observations over a seven-day period of floral visitors to a plot with six flowering plants recorded ten visitors entering the flowers, all representing a species of a mosquito-like insect.

Two specimens were caught on departure from the flowers, preserved and found to be carrying pollen consistent with Thismiaceae. The insects were identified as scuttle flies (Phorideae), probably the genus *Megaselia*, which in other plant groups has a mutualism in which pollination is affected in exchange for the larvae feeding on the decomposing flowers (Sakai 2002; Hall & Brown 1993). It might be expected that other species will have different pollinators since they have pollination structures not seen in *A. kupensis*, such as brightly coloured yellow flowers (vs dull purple and white) and long filiform antennae-like perianth lobes (vs flat, triangular) postulated to disperse pollinator attractant volatiles (Merckx 2008).

Within *Thismia*, “trap” flowers have been suggested, as pollinators have been observed temporarily restrained inside the hypanthium chamber (Guo *et al*. 2019; Nuraliev *et al*. 2021). Self-fertilization could also occur widely in Dioscoreales, including both *Thismia* and *Afrothismia*. The unusual flower structure of *Afrothismia*, with anther appendages extending to the stigma surface at anthesis, indicates this possibility. In contrast, self-fertilzation is strongly indicated by our anatomical sections of *Haplothismia exannulata* (Fig. 3 K, L) showing a mucilagenous mass of germinating pollen tubes growing directly into the style between the carpel margins.

### Seed structure and seed dispersal

Rübsamen (1986) described seeds of two species of *Afrothismia, A.winkleri* (based on *Zenker* 3613, B) and *A. “arachnites”* (based on *Peter s.n*., B). The seeds of the two *Afrothismia* species are 0.7–0.91 x 0.17–0.23 mm, twice the length of the range of the six *Thismia* species described by Rübsamen (1986). She characterised the seeds as elongated, 3–5 cells along the longitudinal axis, and having rows of epidermal cells that are twisted either clockwise or counter-clockwise. The epidermal cells are elongate with raised anticlinal walls showing a suture. The outer periclinal wall is smooth to verrucose, usually collapsed, showing the parallel-bar-like thickenings on the inner periclinal wall (Rübsamen 1986: tafel 82 f–h). Rübsamen (1986) neither discussed nor depicted terminal seed appendages (Fig. 2C).

Burmanniaceae and Thismiaceae are usually classified as having “dust seeds”, which have the possibility of being dispersed by the wind, although in forest floor habitats, even breezes are usually absent, so Maas *et al*. (1986) considered that they are more likely to be dispersed in runnels of water. However, *Afrothismia* seeds were reported by Rübsamen (1986) to be about twice as large as those of a sample of six species of *Thismia*, reaching 0.7–0.91 mm long in *A. winkleri* and *A. “arachnites”*. Measurements of *Afrothismia hydra* at 0.7–0.8 mm long also fit this range (Sainge & Franke 2005). However, seeds of *A. zambesiaca* were recorded at 0.5 mm long Cheek 2009), and those of *A. foertheriana* at 0.5–0.6 mm long (Franke *et al*. 2004), yet those of *A. kupensis* are c. 1.5 mm long (Cheek *et al*. 2019). For other species, seed dimensions are not recorded. In both *A. zambesiaca* (Cheek 2009) and *A. “arachnites”* (Cheek, pers. obs. 2022) the base and apex of the seed have attached white bodies, possibly elaiosomes (Fig. 2C), which suggests that ant dispersal might be possible, rather than the “splash-cup” mechanism believed to occur in *Thismia*. Since the seeds are elevated above the fruit on a “placentophore” in all known *Afrothismia* species where fruits have been observed, the splash-cup mechanism suggested for *Thismia* (Stone 1980) cannot occur in *Afrothismia*.

## Discussion & Conclusions

Setting aside the seven undescribed species listed above, *Afrothismia* (Afrothismiaceae) with 16 accepted species, are the most species-diverse fully mycotrophic genus and family in Africa. In recent years many facets of the biology of this fascinating group have been uncovered, yet important aspects remain poorly known, such as pollination biology, or entirely unreported, such as microsporogenesis, cytology, and seed dispersal, and, we do not yet know which autotrophic plant species their soil fungal symbionts depend upon.

However, the highest priority for the family must be to protect from extinction the known species, to conserve them in their natural habitats e.g. by including them in Important Plant Areas (Darbyshire et al. 2017) and developing species conservation action plans to improve the likelihood of their survival (e.g. Couch *et al*. 2022). This is crucial since *ex situ* conservation is currently not possible. For its size, the genus must be amongst the most highly threatened on the planet, because 11 of the 14 assessed species are globally Critically Endangered, the highest level of threat. Although only one of these species is considered extinct, several others have not been seen alive in decades. Searching for these long-lost species is urgent. *Afrothismia* accepted species numbers have increased in the last 20 years by 300%, from 4 to 16. It can be projected that more await discovery so long as unsurveyed suitable habitat survives, and it is vital to find and protect these also.

Until species are documented, described and known to science, it is difficult to assess them for their IUCN conservation status, and therefore the possibility of conserving them is reduced (Cheek *et al*. 2020). Documented extinctions of plant species continue. In the Cross-Sanaga Interval of Cameroon, the centre of diversity for *Afrothismia* with half the accepted species, the best documented global species extinction is another fully mycoheterotrophic species, *Oxygyne trianda* Schltr. (Thismiaceae, Cheek & Onana 2011; Onana & Cheek 2011; Cheek *et al*. 2018a). Examples of species becoming extinct before they are known to science appear to be on the increase. In Cameroon, inside the Cross-Sanaga Interval, examples are *Vepris bali* Cheek and *Monanthotaxis bali* Cheek (Cheek *et al*. 2018c; Cheek *et al*. 2022). In all cases, anthropogenic habitat clearance for agriculture has been the cause of these extinctions.

## Acknowledgements

The first author thanks R. Vogt (B) for access to material of *Afrothismia ‘arachnites’*, Janis Shillito for typing and Anne Marshall for retrieval of literature.

